# Enhanced Formulation of Precision Probiotics through Active Machine Learning

**DOI:** 10.1101/2025.07.17.665298

**Authors:** Anweshit Panda, Manaswani Adhikari, Sourya Subhra Nasker, Anish K. Nayak, Saroj K. Nayak, Sita. K. Dash, Sasmita Nayak

## Abstract

The human gut microbiome is crucial to health, with dysbiosis increasingly linked to disease. Precision probiotics offer a promising approach to restoring microbial balance, but ensuring probiotic viability through gastrointestinal transit remains a challenge. This study applies an advanced active machine learning (ML) approach to predict how excipients affect the growth of *Lactobacillus plantarum*, a commonly used probiotic. State-of-the-art experiments were carried out to complement the ML study. Starting with five known excipient- probiotic interactions, we apply active ML over three rounds to predict the effects of 116 excipients, iteratively refining model certainty and accuracy. Five ML models—Neural Networks, Gradient Boosting, Logistic Regression, Random Forest, and Support Vector Machines—were trained and evaluated, with the final model achieving certainty levels close to 90%. Unlike previous methods, which retrained new models per iteration, our approach continuously optimized a single model, enhancing prediction stability and reducing error. These results highlight the potential of active ML to support accurate excipient selection in probiotic formulations.

## 1. Introduction

Gut microbiota, also known as the gut flora, encompasses a vast array of microorganisms residing in the gastrointestinal tract (GI tract) of humans. This complex ecosystem comprises bacteria, viruses, fungi, and other microorganisms vital for human health and well-being. The gut microbiota contributes to the prevention and management of various diseases such as inflammatory bowel disease (IBD), irritable bowel syndrome (IBS), metabolic syndrome, neurological disorders, obesity, diabetes, and cancer[1, 2]. Microbial imbalance (also known as dysbiosis) occurs due to various factors such as antimicrobial exposure, low fiber diet, and polypharmacy and is often associated with increased health risk[3, 4].

Probiotics are formulations of live microorganisms that are fundamental in maintaining a healthy gut environment. They modify the gut microbiota composition by simultaneously increasing the number of beneficial microorganisms, suppressing the growth of pathogens and maintaining a balanced body system[5, 6]. Probiotics secrete antimicrobial compounds such as bacteriocins and hydrogen peroxide, which prevent the growth of pathogens and maintain a healthy gut microbiota. They also play a significant role in drug development, offering a new frontier for discovering novel therapeutic agents and approaches.

In response to these findings, a new class of therapeutics, known as microbiome medicines, has been developed with probiotics as a key component to prevent and treat diseases by addressing the dysbiosis [7, 8],. The scientific community is increasingly focusing on defined probiotic strains as potential drug products due to their characterised mechanisms and demonstrated clinical benefits [9–11]. Thus precise identification of next-generation probiotics could revolutionise disease management strategies in the coming years[12–15].

The exact formulation of probiotics is a crucial step for their incorporation into approved medicinal products. Currently, probiotic colonisation of the GI tract is dependent on the type of strains used, and its effects are mostly transient, with variations observed in individuals possessing differing microbiome compositions prior to treatment. Exposure to harsh conditions induced by gastric acid and bile salts in the upper GI tract, can lower probiotic colonisation by reducing the number of viable organisms reaching the distal gut[16, 17]. Strategies to overcome this include using acid-tolerant probiotics and/or protective probiotics. The latter strategy can employ coating technologies that release therapeutics specifically within the colon where microbiota density is highest[18–20]. Delivery to the intended site of colonisation, however, does not guarantee that probiotic strains will outcompete the existing microbiome for attachment[21].

During the development of a probiotic formulation, excipients should be chosen carefully since it can significantly affect the growth and viability of the probiotic. Choosing the right excipients for a quick and effective formulation of probiotics poses a challenge. Previous studies have highlighted the significance of choosing functional excipients, as they can either enhance or inhibit microbial growth, thereby impacting the overall therapeutic effectiveness of the probiotic formulation.

Recently, McCoubrey et al. studied the effects of six excipients (D-mannitol, polysorbate 80, aspartame, guar, β-carotene, and sucrose) on the growth of *Lactobacillus paracasei* [22]. They utilized an active machine learning approach to predict the effects of 111 excipients, achieving an average certainty of 67.70% in their final model. Model certainty refers to the confidence of the machine learning model in its predictions, which is quantified based on the probabilities assigned to each prediction. In contrast, model accuracy reflects the proportion of experimentally validated predictions as correct. The McCoubrey study reported a model accuracy of 75%, indicating that 3 out of 4 predictions were experimentally confirmed to be correct. In contrast, model certainty offers a probabilistic measure of the reliability of predictions across the dataset, and this approach could potentially be refined for probiotic formulation(s).

In this study, we apply an advanced form of machine learning (ML), known as active ML, to analyse and predict excipient effects on the growth of *Lactobacillus plantarum*, enabling a strategic approach towards efficient probiotic microdelivery. This is one of the first comprehensive studies to apply active ML to next-generation probiotics and, more broadly, to microbiome science as a whole[23, 24] . Active ML is well-suited to generating predictions from small datasets where collection of large data is not feasible or desired[25]. During active ML, an initial ML model is developed on available labelled data. The model can then output predictions for unlabelled data, indicating its certainty for each prediction. Users can experimentally evaluate datapoints for which the model is most uncertain and subsequently teach the model the new results, to improve overall model certainty[26].

In contrast to earlier publication, which utilized the random forest method, this study employs various methods such as SVM, logistic regression, gradient boost, and neural networks to predict the effects of 111 pharmaceutical excipients on the growth of a commercially available probiotic, *Lactobacillus plantarum*[27]. Predictions were based on obtaining correlations from a dataset of five excipient-probiotic interactions, which were experimentally evaluated specifically for this project while carrying out routine formulation development. *L. plantarum* is a well-researched probiotic species, with documented benefits for a myriad of diseases including irritable bowel syndrome, cardiovascular disease, and immune system regulation[28–30]. The ML model developed in this study is designed to facilitate the selection of functional excipients for the optimisation of *L. plantarum* proliferation *in vivo*. The methods used can be easily translated for developing precision probiotic, thereby highlighting active ML as a powerful tool for optimising the *in vivo* performance of pharmaceutical formulations.

## 2. Materials and Methods

### 2.1. Probiotic Growth and Excipient Effects

#### 2.1.1 Materials

*Lactobacillus plantarum* was procured from ATCC (American Type Culture Collection, The Global Bioresearch Centre). The probiotic strains were cultivated in de Man, Rogosa, and Sharpe (MRS) media (HiMedia, Cat. no.- M1164) under oxygen-free conditions. Several excipients were sourced from HiMedia including, sodium chloride (Cat. No.-MB023), zinc chloride (Cat. No.- MB046), potassium bicarbonate (Cat. no.- GRM1789), and sucrose (Cat. no.- MB025). Sigma Aldrich provided urea (Cat. No.- U5378) and magnesium sulphate (Cat. No.- M7506). Dextrose (Cat. No. 51758), L-ascorbic acid (Cat. No. 23006), and glycine (Cat. No. 71943) were supplied by SRL, India.

#### 2.1.2 Analysis of L. plantarum Growth

*Lactobacillus plantarum* glycerol stocks were inoculated into MRS media and incubated overnight at 37℃ under anaerobic conditions. The following day, a subculture was performed with a 1:100 dilution, and the cells were again incubated at 37℃ in anaerobic conditions. To analyze the bacterial growth curve, the growth of *L. plantarum* was examined over a period of 48 hours. Hourly bacterial samples were collected for 48 hours, and their optical density (OD) was measured at 600 nm[31].

#### 2.1.3 Measuring Excipient Effects

Excipients were selected and added to MRS broth to assess their role in the growth of *Lactobacillus plantarum.* Each investigated excipient (sodium chloride, zinc chloride, potassium bicarbonate, sucrose, urea, magnesium sulphate, dextrose, L-ascorbic acid, glycine) was incorporated separately into the MRS broth at a concentration of 70 µM. This concentration was chosen because drugs are generally found at levels of 20 µM in the terminal ileum and colon, where microbiota populations are most abundant[32]. The growth of *L. plantarum* in presence of 70 µM excipients were analysed from optical density (OD) values recorded at 600 nm. The results were plotted from three (n=3 for each excipient) independent sets of experiments. Growth curves obtained in the presence of excipients were compared with growth curves obtained in the pure microbial growth medium (Figure S1-S10, Supplementary Materials). The area under the curve (AUC) and time taken for maximum growth (T_max_) were used as comparative features[22]. The AUC values were obtained using the equation:

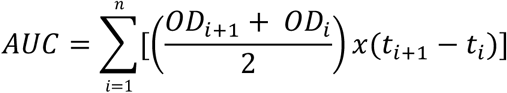

where OD_i_ is the measured optical density. The results were analyzed using online DMFit from ComBase software (www.combase.cc), and the T_max_ (time taken for maximum growth) values were obtained. A one-way analysis of variance (ANOVA) was performed using the GraphPad Prism version 5.01 for Windows, GraphPad Software, San Diego, CA, U.S.A. (www.graphpad.com). This was used to assess whether there were significant differences (p < 0.05) between microbial growth in the presence of excipients, compared with the absence of excipients (growth media only). The excipient was determined to be inhibiting microbial growth if AUC was significantly reduced or there was an increase in T_max_. Conversely, if an excipient significantly increased AUC, or decreased T_max_, then the excipient was said to be promoting microbial growth. If excipients did not significantly alter AUC or T_max_ they were viewed as neutral; neither inhibiting nor promoting microbial growth. The initial dataset consisted of nine excipients—sodium chloride, zinc chloride, potassium bicarbonate, sucrose, urea, magnesium sulphate, dextrose, L-ascorbic acid, and glycine— chosen to represent a diverse array of chemical and functional characteristics relevant to probiotic development [22].

### 2.2. Machine Learning Approaches

#### 2.2.1 Dataset Curation

The initial dataset was carefully curated to include five excipients with distinct effects on *Lactobacillus plantarum* growth: doxycycline hydrochloride, L-ascorbic acid, glycine, dextrose, and zinc chloride. These compounds were selected to provide a balanced representation of effects, with two growth promoters (L-ascorbic acid, dextrose), two growth inhibitors (doxycycline hydrochloride, glycine), and one neutral compound (zinc chloride) [22]. This balanced distribution allowed for the development of preliminary models with robust predictive capacity across excipient-response relationships.

For active learning, three additional compounds—magnesium sulphate, sucrose, and urea were selected and tested iteratively over three rounds to refine model predictions. Each excipient was confirmed to be soluble in the aqueous bacterial growth medium, ensuring effective interaction with *L. plantarum*. The excipients selected for both the initial dataset and the active learning pool spanned a wide chemical and functional space, capturing a diverse range of potential interactions. For final validation, two neutral compounds, sodium chloride and potassium bicarbonate, were included to assess model performance on previously untested neutral effects.

This careful and strategic dataset curation ensures the comprehensive exploration of excipient effects on *L. plantarum* growth. Full details of the labelled dataset and active learning pool are provided in the Supplementary Materials to support further studies.

#### 2.2.2 Feature Selection and Preprocessing

In this study, a comprehensive set of approximately 1600 chemical descriptors were associated with each excipient in both the labelled datasets and the unlabelled pool. These descriptors were derived from each excipient’s simplified molecular-input line-entry system (SMILES) structure of each excipient, sourced from the PubChem database[33] . To transform the SMILES annotations of the excipients into a robust set of molecular features, Mordred, a state-of-the-art molecular descriptor calculator was deployed[34]. This transformation provided a diverse array of molecular features, encompassing both computational descriptors and more traditionally recognised chemical features. Features such as ‘4-membered aromatic ring count’ and ‘number of atoms’, allowed for a rich, multi-dimensional representation of each excipient. To ensure a fair and unbiased machine learning process, all features underwent a scaling process using Python’s Standard Scaler. This crucial preprocessing step normalised the range of the descriptor values, thereby eliminating any potential bias due to the different units of the features and allowing each feature to contribute equally to the machine learning models.

#### 2.2.3 Data Visualisation

To visualize the relationships between the labelled excipients, we employed Principal Component Analysis (PCA), a widely used technique for reducing the dimensionality of high-dimensional datasets. PCA works by identifying new axes, called principal components, which are linear combinations of the original features and are orthogonal to each other. These components are ranked based on the proportion of variance they explain. The variance in PCA refers to the spread or variability of the data in the original dataset across its features. Each principal component explains a portion of this variability, with the first principal component capturing the direction in which the data points differ the most. Subsequent components capture remaining variability in decreasing order, ensuring the main patterns in the dataset are prioritized. Utilizing the PCA functionality within Python’s scikit-learn package (version 0.23.2), we transformed our multidimensional feature space into a two-dimensional space, facilitating easy visualization and interpretation[35].

In our analysis, we plotted the first principal component (PC1) against the second principal component (PC2) to capture the most significant relationships in our data on a two-dimensional graph. The transformation focuses on retaining the most important patterns in the data, which are associated with the largest sources of variation. By discarding less significant patterns—those that contribute minimally to the variation—the transformation reduces the complexity of the dataset while preserving the main trends or groupings that are present in the data. Upon visually inspecting the resulting PCA plots, patterns and clusters among the labelled excipients were observed. An effort was made to identify instances where excipients with the same label grouped together in the feature space. This indicated similar effects on *L. plantarum* growth. This visualization provided insights into the grouping tendencies of excipients and their potential roles in influencing microbial growth.

### 2.3. Active Machine Learning

In this study, we employed the Active Learner and Uncertainty Sampling packages of the modAL active learning framework for Python (version 0.4.1) to construct our machine learning pipeline. The initial machine learning model considered was the Random Forest classifier. Random Forest is an ensemble learning algorithm that builds multiple decision trees during training and combines their outputs to improve prediction accuracy and reduce overfitting[25, 36]. A key configurable parameter in Random Forest is the random state, which controls the randomness of the algorithm, ensuring reproducibility of results. The first step involved fitting the initial labelled dataset to a Random Forest Classifier from Python’s scikit-learn package, with the random state set to 0. In this study, the random state was set to 0 to ensure reproducibility by controlling randomness in processes such as data sampling and tree construction, which impacts the overall consistency of the model. This base model was selected due to its computational efficiency and its ability to interpret nonlinear data relationships, while simultaneously limiting overfitting.

The ActiveLearner class from the modAL framework was utilized to integrate machine learning models into the active learning pipeline. This class provides a framework for iterative model training and querying, leveraging uncertainty sampling to identify the most informative samples for labeling. By encapsulating each model, including Random Forest, SVM, Logistic Regression, Gradient Boosting, and Neural Networks, the ActiveLearner ensured a consistent and efficient process for dataset augmentation and model refinement.

Utilizing the modAL framework, we computed predictions for how the 116 excipients present in the unlabeled pool would influence the growth of *Lactobacillus plantarum.* Each prediction was assigned an uncertainty score ranging from 0 to 1.00, with higher scores indicating greater uncertainty. To improve model certainty, uncertainty sampling, a commonly used method in active machine learning, was implemented[37]. In uncertainty sampling, an unlabeled excipient is selected for experimental testing based on its relevance or importance as determined by the active learning algorithm. This process of selection is known as querying.

For each query, we selected the unlabeled excipient deemed most relevant to the observed growth rate trends of *Lactobacillus plantarum*. Upon measuring its true label experimentally, it was incorporated into the labelled dataset and taught to the machine learning model. After updating the labelled dataset, the model generated revised predictions and updated uncertainty scores for the remaining unlabeled excipients. Further queries were sequentially made using this method to actively enhance the overall certainty of the machine learning model by experimentally reinforcing the training set.

In this study, a diverse set of machine learning models was employed within the active learning framework to classify the effects of pharmaceutical excipients on the growth of *Lactobacillus plantarum*. These models included Random Forest (as used in the reference study for its computational efficiency and ability to handle non-linear relationships), Support Vector Machines (SVM), Logistic Regression, Gradient Boosting, and Neural Networks. Each model was integrated into the active learning pipeline, enabling iterative improvement of predictions through uncertainty sampling and successive querying of the most informative data points. The use of multiple models provided a comparative perspective on the robustness and effectiveness of different algorithms in identifying excipient effects.

#### 2.3.1 Support Vector Machines (SVM)

Support Vector Machines (SVM) are a powerful and flexible class of supervised learning algorithms used for both classification and regression tasks[38]. SVM works on the principle of identifying the appropriate hyperplane in the feature space for separating different classes of data points. A hyperplane is a decision boundary that partitions the feature space into regions, each corresponding to a specific class. The SVM algorithm identifies the hyperplane that achieves the maximum margin—the greatest distance between the hyperplane and the nearest data points from any class, known as support vectors. This margin maximization reduces overfitting and improves the generalization of the model to unseen data[39, 40].

For our active learning framework, the SVM model was implemented using the SVC class from the scikit-learn library in Python. We configured the SVM with a linear kernel, which assumes the data to be linearly separable (i.e., the classes can be separated by a straight line or flat hyperplane in the feature space). Additionally, probability estimates were enabled, allowing the SVM to assign probabilistic scores to its predictions. These scores facilitated uncertainty sampling by quantifying the confidence in the model’s predictions, a key component of the active learning process.

The active learning process utilized the ActiveLearner class, initialized with the customized SVM estimator, to iteratively select and label samples predicted to be the most informative based on their uncertainty scores. Model evaluation focused on uncertainty-based metrics, such as model certainty and error bars, to dynamically assess the SVM’s predictive accuracy and robustness[41]. A more detailed explanation of the methodology and additional performance analyses is included in the supplementary material.

#### 2.3.2 Logistic Regression

Logistic Regression is a statistical model used for binary classification tasks, where a logistic function maps input features to probabilities. This allows the model to classify data into two categories. This model is particularly effective for problems where the relationship between features and the target variable is approximately linear[42]. In this study, Logistic Regression was integrated into the active learning framework to classify the effects of pharmaceutical excipients on the growth of *Lactobacillus plantarum*. The model was implemented using the LogisticRegression class from Python’s scikit-learn library[43].

This active learning framework also employed uncertainty sampling, a strategy that prioritizes samples for which the model is least confident. For Logistic Regression, high uncertainty samples were chosen by focusing on data points with prediction probabilities closest to 0.5. These uncertain samples were labeled and added to the training dataset, allowing the model to iteratively refine its decision boundary and improve its performance.

Machine learning models such as Logistic Regression have hyperparameters that can be fine-tuned for optimal performance. Default hyperparameter settings were used for Logistic Regression without additional tuning. Further optimization was deemed unnecessary as high performance was already achieved by the active learning pipeline with the current configurations[44]. The model’s performance was evaluated using uncertainty-based metrics such as model certainty and error bars, which assessed predictive reliability across successive active learning cycles[45]. Detailed explanations of the logistic function, decision boundary, uncertainty sampling, and evaluation metrics are provided in the supplementary material.

#### 2.3.3 Gradient Boosting

Gradient Boosting is an ensemble learning technique that constructs a strong prediction model by combining multiple weak learners, typically decision trees[46]. The model is built in a stage-wise manner, where each stage aims to minimize a loss function that measures the difference between predicted and actual values. The loss function, which quantifies the model’s prediction errors, is a key component in guiding the iterative optimization process. In Gradient Boosting, the differentiable nature of the loss function allows the algorithm to compute gradients, which are used to iteratively reduce errors and improve predictive accuracy.

In this study, Gradient Boosting was implemented using the GradientBoostingClassifier class from the scikit-learn library in Python[43]. The model was integrated into the active learning framework via the ActiveLearner class, enabling iterative training and querying. The active learning process employed uncertainty sampling to select the most informative samples for labeling, focusing on excipients where the model exhibited the highest uncertainty in its predictions.

Default hyperparameter settings were used for the Gradient Boosting model without additional tuning. This decision was based on the sufficient performance observed within the active learning framework[44]. Model performance was evaluated using uncertainty-based metrics, such as model certainty and error bars, to dynamically assess improvements in predictive reliability across successive active learning cycles[45]. Detailed explanations of the Gradient Boosting algorithm, loss function, and evaluation methodology are provided in the supplementary material.

#### 2.3.4 Neural Networks

Neural Networks are a set of algorithms modeled loosely after the human brain, designed to recognize patterns in data. They achieve this through a layered architecture, where data flows through interconnected layers of artificial neurons[47]. In this study, a deep neural network was implemented using the Keras library with a Sequential architecture to classify the effects of pharmaceutical excipients on the growth of *Lactobacillus plantarum*.

The neural network consisted of two fully connected Dense layers, with 64 and 32 neurons, respectively, followed by Dropout layers to mitigate overfitting by randomly disabling neurons during training. The output layer contained three neurons with a softmax activation function, which transformed raw outputs into probabilities corresponding to the three target classes. The model was compiled using the Adam optimizer and the categorical cross-entropy loss function, with accuracy set as the primary evaluation metric. Training was conducted over 50 epochs with a batch size of 32 to ensure sufficient learning from the initial labeled dataset.

To integrate the neural network into the active learning framework, a custom KerasWrapper class was implemented. This wrapper enabled compatibility with scikit-learn’s ActiveLearner class, facilitating iterative training, querying, and prediction within the active learning process. The active learner employed an uncertainty sampling strategy to identify and label the most informative samples, thereby enhancing model performance[44]. Further explanations of neural network architecture, key hyperparameters, and training concepts are provided in the supplementary material.

## 3. Results and Discussion

### 3.1. Effect of Excipients on Bacterial Growth

Nine excipients were chosen for evaluating their effect on the growth of *L. plantarum,* and the results are shown in Table 1. It was found that 6 of the 9 excipients had no effect on bacterial growth. L-ascorbic acid was observed to promote growth while glycine impaired growth. Dextrose was found out to be partially promoting.

**Table 1.** Effects of excipients on the growth of *L. plantarum*. Here, AUC: area under curve; T_max_: time taken for bacteria to reach maximum growth. The statistical significance (p-value) between the control and experimental groups is denoted by ns (p > 0.05), or * (p ≤ 0.05), or ** (p ≤ 0.01), or *** (p ≤ 0.001), or **** (p ≤ 0.0001).

| Excipients | Observed effect | P-value<br>(AUC and T<sub>max</sub>) | Label |
| --- | --- | --- | --- |
| Sodium Chloride | No change | 0.1 (ns), 0.97 (ns) | Neutral |
| Urea | No change | 0.93 (ns), 0.87 (ns) | Neutral |
| Magnesium Sulphate | No change | 0.64 (ns), 0.31 (ns) | Neutral |
| Sucrose | No change | 0.96 (ns), 0.1 (ns) | Neutral |
| Potassium bicarbonate | No change | 0.1 (ns), 0.99 (ns) | Neutral |
| Zinc chloride | No change | 0.1 (ns), 0.93 (ns) | Neutral |
| Glycine | AUC decreased, T<sub>max</sub> increased | 0.04 (*), 0.025 (*) | Inhibit |
| Dextrose | AUC increased, T<sub>max</sub> decreased | 0.01 (*), 0.1 (ns) | Promote |
| L-ascorbic acid | AUC increased, T<sub>max</sub> decreased | 0.03 (*), 0.025 (*) | Promote |

L-ascorbic acid, a multifunctional compound, acts as a cofactor, enzyme complement, co-substrate, and potent antioxidant in various metabolic processes. It also helps stabilize vitamin E and folic acid while boosting iron absorption [48]. In the presence of L-ascorbic acid, the growth of *L. plantarum* was accelerated (AUC increased, p = 0.03 and T_max_ decreased, p = 0.026). Glycine is a crucial non-essential amino acid primarily synthesised in kidney and liver. However, this is often produced in humans and animals at insufficient amounts under typical feeding conditions. It serves as a precursor for important metabolites and has been shown to improve health, support development, and aid in the prevention and treatment of various diseases, including metabolic disorders, cardiovascular issues, and cancer[49]. In the current study, exposure to glycine, however, reduced the growth of *L. plantarum* (AUC decreased, p = 0.038, and T_max_ increased, p = 0.026). Dextrose promoted the growth of *L. plantarum*, as indicated by a significant increase in AUC (p = 0.01). However, it did not accelerate the growth rate of the probiotic bacteria in the study, as indicated by a decrease in T_max_ (p = 0.1, ns). This indicates that the effectiveness of dextrose as a carbon source is strain-dependent across probiotics. Therefore, the variation in excipient activity may reflect their specific role(s) under varied physiological conditions, and heterogeneous effects among different strains of *Lactobacilli*. For instance, a 2018 study found sodium chloride to be an inhibitory excipient that significantly reduces the growth rate of *L. plantarum* ATCC 8014, with a negative correlation observed between NaCl concentrations and microbial growth under the tested conditions[50]. Similarly, a 2022 study reported that *L. paracasei* exhibited neutral effects when paired with excipients such as sucrose, mannitol, and polysorbate 80, whereas β-carotene enhanced its activity and guar acted as an inhibitor[22]. Likewise, prior and recent literature further underscores the experimental evidence of heterogeneity in activity of excipient (selected for this study) towards different probiotic strains[51–56].

The following section of the manuscript will give a detailed explanation of the strategies employed through active machine learning (ML) to analyze the observed effects on accuracy prediction. These methodologies highlight the intricate relationships within the data and demonstrate how active ML can optimize predictive performance by iteratively refining models based on real-time feedback. By leveraging these advanced techniques, we show the impact on enhancing prediction accuracy and overall model efficacy.

### 3.2. Data interpretation using Principal Component Analysis

As mentioned earlier, Principal Component Analysis (PCA) was utilized to explore the underlying structure of the dataset at different stages of the active learning process. This dimensionality reduction technique provided a visual representation of how the data distribution evolved from the initial phase of active learning to the final query, offering insights into the effectiveness of the active learner in capturing the most informative samples.

#### 3.2.1. Data Evolution during Active Learning

At the commencement of the active learning process, the dataset comprised a limited and potentially non-representative subset of the entire feature space. Applying PCA to this initial dataset (Figure 2A) revealed a concentrated clustering of data points, indicating that the selected samples were insufficient to capture the inherent diversity of the dataset fully. This phase laid the foundation for the active learner to identify regions of uncertainty or high information density.

**Figure 1.**
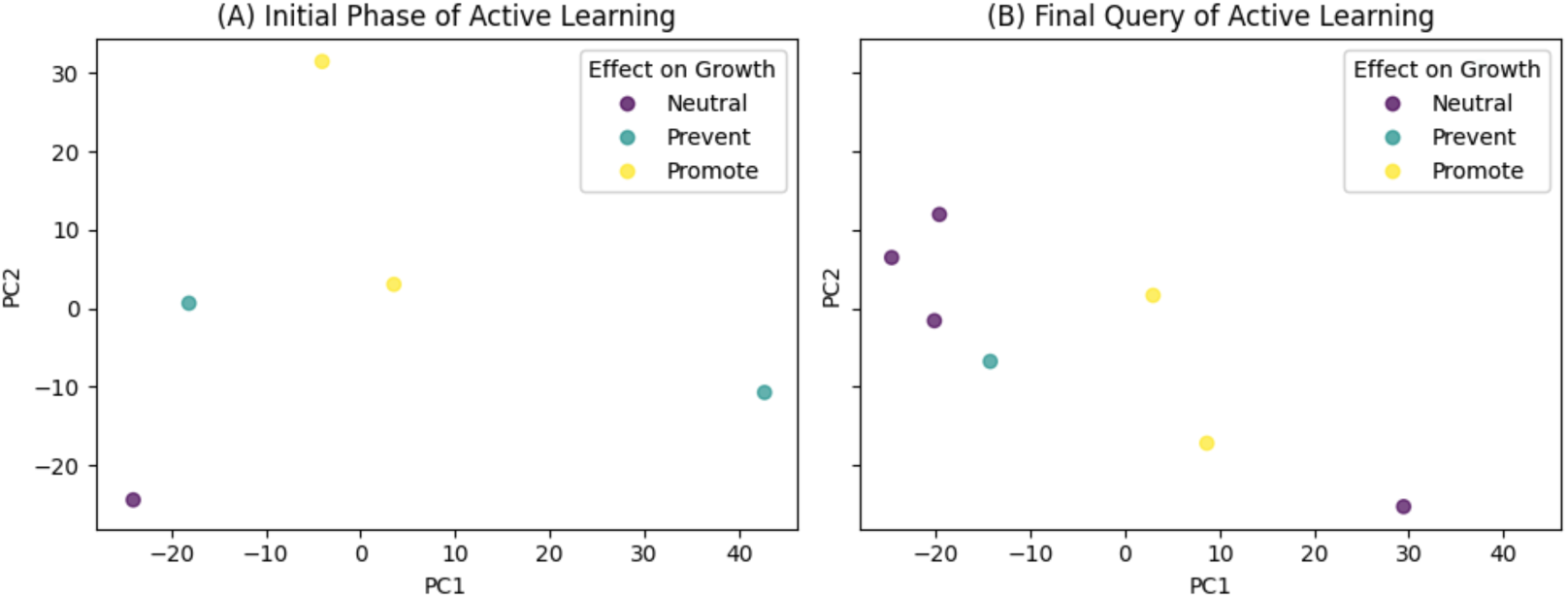
Principal Component Analysis (PCA) of the Dataset During Active Learning. (A) Initial Phase of Active Learning: This subplot illustrates the PCA results during the initial phase of active learning, where data points are concentrated in a small region of the feature space. The clustering reflects the limited diversity of the initial dataset, potentially constraining the model’s ability to generalize. Different classes (’Neutral’, ’Prevent’, ’Promote’) are color-coded to highlight the initial class distribution and the compactness of feature representations. (B) Final Query of Active Learning: This subplot presents the PCA results after the final query of active learning. Three additional excipients were added to the initial dataset of five, resulting in a more dispersed data distribution within the feature space. This dispersion demonstrates the incorporation of diverse and informative samples, reflecting the active learning process’s effectiveness in expanding the dataset’s feature representations. The updated class distribution highlights the model’s improved ability to capture diverse data characteristics, thereby supporting better generalization and performance.

**Figure 2.**
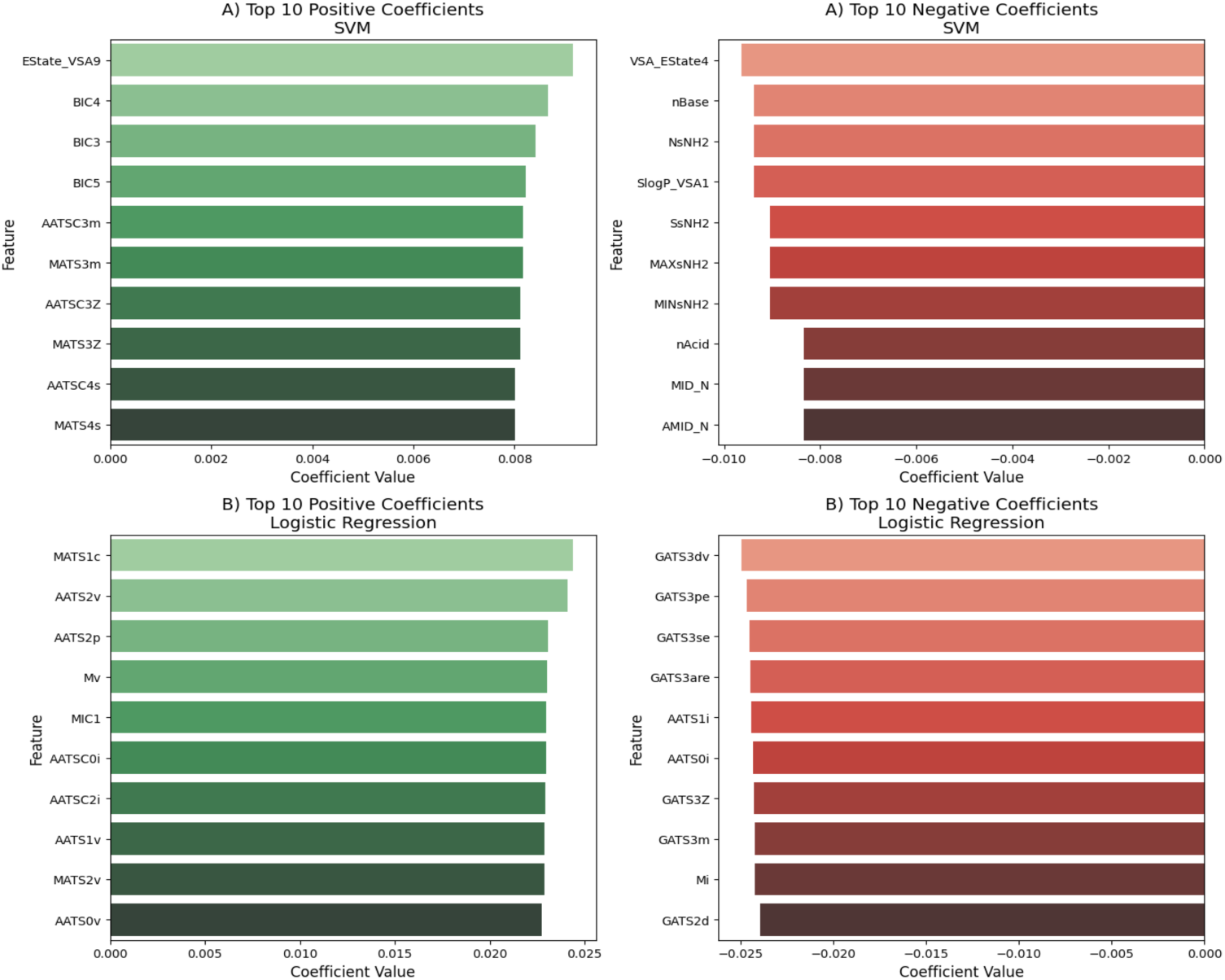
Correlation of Feature Coefficients for Linear Models. (A) Support Vector Machine (SVM): This subplot illustrates the correlation of feature coefficients in the SVM model. Positive coefficients, such as those of *EState_VSA9* **(0.00915)** and *BIC4* **(0.00866)**, indicate that an increase in the feature value enhances the likelihood of the positive class outcome. Conversely, negative coefficients, including *VSA_EState4* **(-0.00963)** and *nBase* **(- 0.00938)**, signify that higher values of these features are associated with a decreased probability of the positive class. These bidirectional effects emphasize the nuanced roles that different features play in shaping the decision boundary of the SVM model, with some features contributing positively and others negatively to the classification task. **(B) Logistic Regression:** This subplot presents the correlation of feature coefficients in the Logistic Regression model. Positive coefficients, such as those for *MATS1c* **(0.02438)** and *AATS2v* **(0.02409)**, contribute to increasing the probability of the positive class, while negative coefficients, such as *GATS3dv* **(-0.02495)** and *GATS3pe* **(-0.02466)**, reduce the likelihood of the positive class outcome. These relationships provide valuable insights into how individual features directly influence the model’s predictions, offering a deeper understanding of their roles in the classification process.

As active learning progressed, the algorithm iteratively selected and queried new samples expected to provide the most significant information gain. Upon reaching the end of the active learning cycle, PCA was carried out again on the augmented dataset (Figure 2B). The resulting plot demonstrated a more dispersed and comprehensive distribution of data points across the principal components. This dispersion highlighted the active learner’s ability to identify and incorporate diverse, informative samples, thereby enhancing the model’s generalization capability across the feature space.

### 3.3. Correlation of Features in Different Machine Learning Algorithms

#### 3.3.1. Feature Importance in Linear Models

Understanding the correlation between features is crucial for interpreting model behaviour and identifying potential multicollinearity issues. Multicollinearity occurs when two or more features are highly correlated, which can lead to unreliable and unstable estimates of the model coefficients. This makes it difficult to determine the individual effect of each feature on the target variable. In linear models like Support Vector Machines (SVM) and Logistic Regression, feature coefficients offer insight into how each feature relates to the target variable. A positive coefficient indicates that as the feature value increases, the likelihood of the positive class also increases, while a negative coefficient suggests the opposite. The magnitude of the coefficient reflects the strength of this relationship.

##### Support Vector Machine (SVM)

The correlation plot for SVM (Figure 3A) highlights both positive and negative associations between features and the target class. For instance, features like *EState_VSA9* (0.00915) and *BIC4* (0.00866) have positive coefficients, indicating that increases in these feature values enhance the likelihood of a positive class outcome. In the context of bacterial growth, these features may be linked to molecular or environmental factors that promote bacterial proliferation, offering insights into underlying biological mechanisms. Conversely, features such as *VSA_EState4* (-0.00963) and *nBase* (-0.00938) exhibit negative coefficients, suggesting that higher values of these features decrease the probability of the positive class, potentially reflecting characteristics that inhibit bacterial growth. These opposing correlations provide a nuanced understanding of how specific features influence the SVM model’s decision boundary and contribute to predictions related to bacterial proliferation.

**Figure 3.**
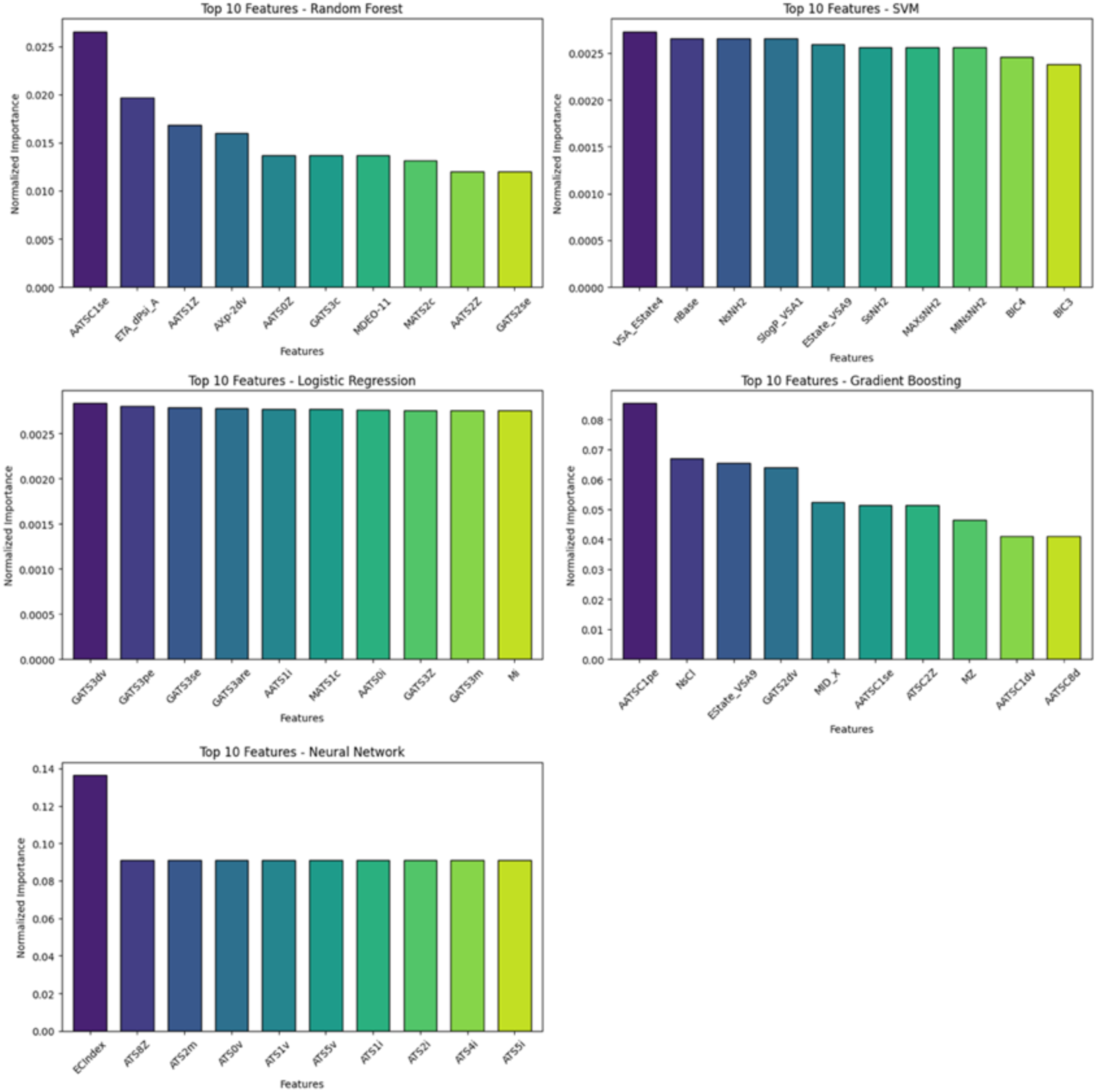
Feature Importance Plots Across Machine Learning Models. This figure presents plots depicting the top ten features for each machine learning model utilized in the study, including Random Forest, Support Vector Machine (SVM), Logistic Regression, Gradient Boosting, and Neural Networks. Feature importance scores are normalized to facilitate comparison across different models, highlighting both common and unique influential features. For instance, *AATSC1se* and *ETA_dPsi_A* are prominent in the Random Forest model, while the Neural Network model emphasizes ECIndex and various ATS descriptors, underscoring each model’s distinct approach to feature significance and predictive performance.

**Figure 4.**
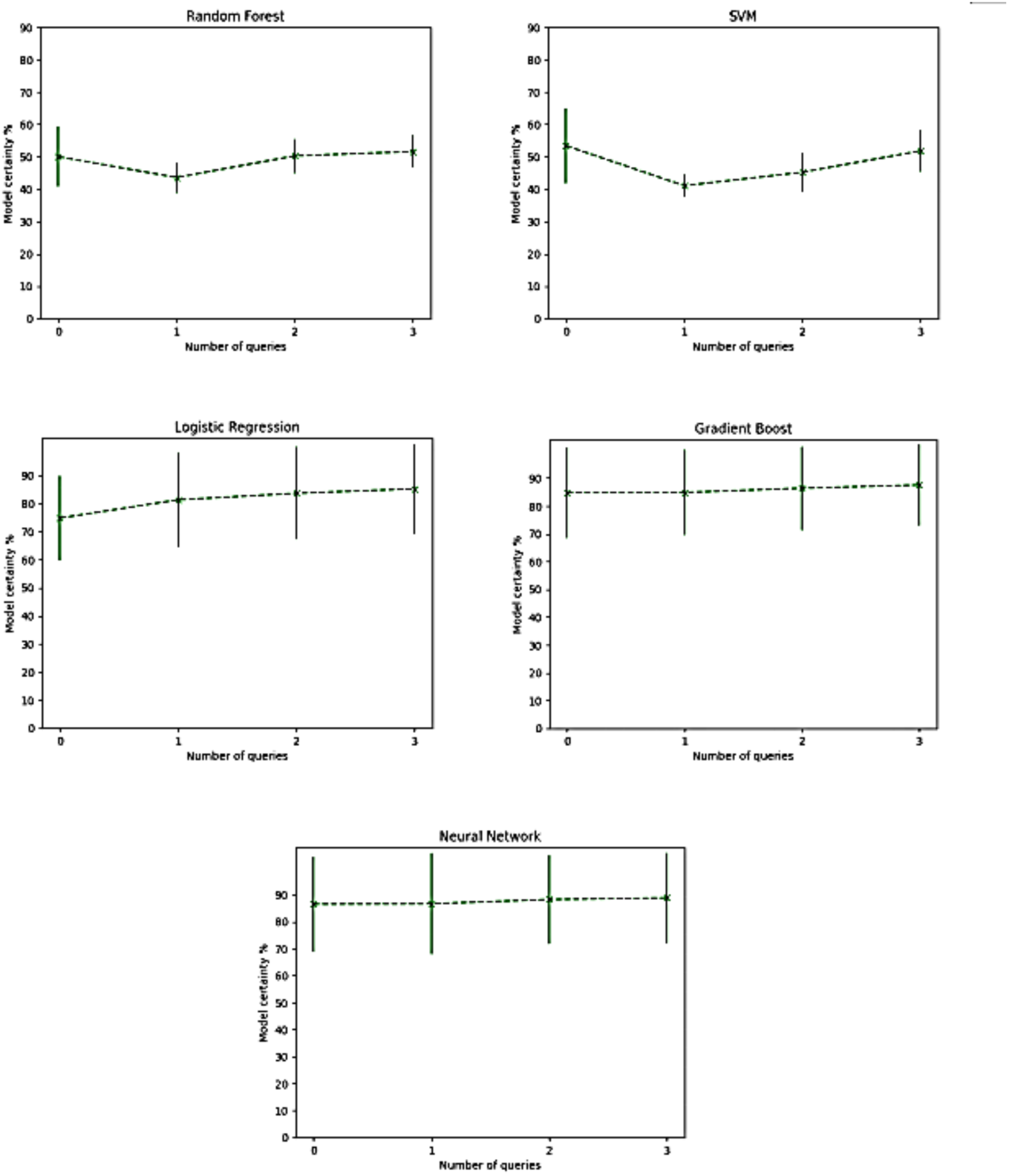
Model Certainties Across Active Learning Queries. The figure showcases the certainty levels achieved by five distinct machine learning models—Neural Networks, Gradient Boosting, Logistic Regression, Support Vector Machines (SVM), and Random Forest—across three rounds of active learning queries. Each model’s performance is depicted through its certainty percentage, illustrating how predictive confidence evolves with each active learning iteration. Notably, Neural Networks and Gradient Boosting exhibit consistent increases in certainty, reaching 88.84% and 87.57% respectively by the final query, demonstrating their robust capacity to enhance predictive stability through iterative learning. Conversely, SVM and Random Forest display lower and more variable certainty levels, highlighting the challenges simpler algorithms face in maintaining consistent performance within complex, non-linear datasets. Logistic Regression maintains steady improvement, achieving 85.33% certainty, which underscores its effectiveness in modelling feature interactions. Overall, the figure emphasizes the superior ability of more complex models to sustain and build upon predictive certainty in active learning scenarios, while simpler models may require additional refinement to achieve comparable stability and accuracy.

##### Logistic Regression

The Logistic Regression correlation plot (Figure 3B) also reveals both positive and negative relationships between features and the target variable. Positive coefficients for features like *MATS1c* (0.02438) and *AATS2v* (0.02409) indicate that increases in these values enhance the likelihood of the positive class. These features may represent molecular properties that are favorable to bacterial growth or survival. On the other hand, features like *GATS3dv* (-0.02495) and *GATS3pe* (-0.02466) possess negative coefficients, suggesting they decrease the probability of the positive class. Biologically, these features may correspond to factors that inhibit bacterial proliferation, such as antimicrobial properties or adverse environmental conditions. These patterns provide critical insights into how individual features contribute to the Logistic Regression model’s predictions and help elucidate the biological drivers of bacterial growth outcomes.

#### 3.3.2. Feature Importance in Tree-based Methods and Neural Networks

Unlike linear models, ensemble tree-based methods such as Random Forest and Gradient Boosting assign feature importance scores that are inherently non-negative. These scores are typically based on metrics like the mean decrease in impurity. Here, impurity measures how mixed the classes are within a node. Standard impurity metrics include Gini impurity and entropy. These quantified the likelihood of incorrect classifications when a randomly chosen element was labelled according to the distribution of labels in the node. Another way of interpreting this would be to view the mean decrease in accuracy, thereby reflecting each feature’s overall contribution to the model’s predictive performance. However, these scores do not indicate whether a feature positively or negatively influences the target variable.

##### Random Forest and Gradient Boosting

In Random Forest and Gradient Boosting models, a feature’s importance is determined by its contribution to improving the model’s accuracy or reducing impurity, regardless of the direction of its effect on the target variable. Consequently, while these methods rank features by their importance, they do not reveal whether a feature increases or decreases the likelihood of the target variable. This contrasts with linear models, where coefficients directly indicate the direction of a feature’s influence.

##### Neural Networks

Neural networks do not provide straightforward feature importance scores due to their complex and non-linear architectures. The interactions between multiple layers and neurons make it challenging to discern the direct influence of individual features on the output. Techniques such as permutation importance or SHAP (Shapley Additive Explanations) can be employed to approximate feature importance in neural networks, but these methods still do not inherently indicate the direction of each feature’s effect.

#### 3.4. Feature Importance

Understanding the relative importance of features in predictive modelling is crucial for interpreting model behaviour, enhancing model performance, and guiding feature selection. In this study, feature importance was evaluated across multiple machine learning models, including Random Forest, Support Vector Machine (SVM), Logistic Regression, Gradient Boosting, and Neural Networks. Each model employs distinct mechanisms to assess feature significance, providing a comprehensive overview of the dataset’s influential variables.

In ensemble tree-based methods like Random Forest and Gradient Boosting, feature importance is calculated based on how much each feature helps to reduce impurity across all trees in the model. A decision node is a point in the tree where the data is split based on a specific feature to create two or more subsets. Impurity measures how mixed or disorganized the data is at a decision node, with common measures including Gini impurity (which reflects the chance of misclassifying a sample) and entropy (which reflects the level of uncertainty). When a feature is used at a decision node to split the data, it reduces impurity by creating subsets that are more uniform. The total reduction in impurity caused by a feature, summed over all trees, determines its importance. This approach identifies features that play a key role in splitting the data effectively, which in turn helps improve the model’s accuracy.

In the case of Support Vector Machine (SVM) and Logistic Regression, feature importance can be inferred from the model coefficients. For a given model, these coefficients (*β_j_*) represent the weight or contribution of each feature (*x_j_*) in the prediction process. In a linear model, the prediction can be expressed as:

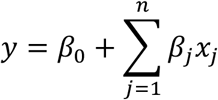

where *y* is the predicted output, *β_0_* is the intercept, and *β_j_* are the coefficients corresponding to each feature *x_j_*. In Logistic Regression, this equation is used as the input to a sigmoid function to calculate the probability of the target class, while in SVM, it helps define the decision boundary. The magnitude of the coefficients (|*β_j_*|) indicates the strength of influence of each feature on the model’s predictions. Larger absolute values suggest that the corresponding feature has a stronger impact on either the position of the decision boundary (in SVM) or the probability of the target class (in Logistic Regression). This makes coefficients a useful way to interpret feature importance in linear models.

For Neural Networks, feature importance was assessed using the theory of permutation importance. This method measures the increase in the model’s prediction error when a feature’s values are randomly shuffled, thereby disrupting the feature’s relationship with the target. This technique captures the contribution of each feature to the model’s overall performance, especially in capturing complex and nonlinear relationships.

The feature importance scores were normalized for comparison across models. Ten of the most influential features of each model are presented in Figure 3 and further discussed in the subsequent sections.

Figure 3 illustrates ten of the most significant features ranked by their normalized importance scores for each model. The analysis reveals both similarities and contrasts in feature significance across the various algorithms. The most influential features in the Random Forest model, the most influential features include *AATSC1se* (0.02653) and *ETA_dPsi_A* (0.01970), indicating strong contributions from these molecular descriptors. These features likely capture critical structural or physicochemical properties influencing the behaviour of the excipient being predicted.

The Support Vector Machine (SVM) model assigns notably lower feature importance scores compared to tree-based models, with *VSA_EState4* (0.00273) leading. This suggests that while SVM can identify significant features, the measure of their relative contributions are more subtle, possibly due to the model’s emphasis on margin maximization rather than impurity reduction.

Similarly, Logistic Regression assigns modest importance scores, with *GATS3dv* (0.00284) and *GATS3pe* (0.00280) at the forefront. These features are indicative of specific atomic triplet descriptors that influence the binary classification outcome, reflecting their role in capturing essential electronic and structural information relevant to the prediction task.

Gradient Boosting exhibits higher feature importance scores, with *AATSC1pe* (0.08538) and *NsCl* (0.06682) being the most significant. The prominence of these features underscores their robust predictive power within the boosting framework, which sequentially focuses on correcting errors from previous iterations.

The Neural Network model displays the highest feature importance scores among all models, with *ECIndex* (0.13636) and a series of ATS descriptors (e.g., ATS8Z, ATS2m) each contributing equally (0.09091). The uniform importance of multiple ATS features suggests that these descriptors collectively encapsulate essential information for the neural network’s predictive performance. This highlights the model’s capacity to leverage complex and nonlinear relationships within the data.

Across the models, certain features consistently emerge as significant. Notably, *AATSC1se* and *AATSC1pe* are prominent in both Random Forest and Gradient Boosting, indicating that these features likely represent key molecular attributes that significantly influence the target variable. Additionally, features such as *GATS3dv*, *GATS3pe*, and *GATS3se* are prominently featured in Logistic Regression, underscoring their role in capturing essential electronic and structural information relevant to the prediction task.

The Neural Network model highlights a range of ATS descriptors (e.g., *ATS8Z*, *ATS2m*, *ATS0v*), which are highly influential and may reflect complex interactions and nonlinear relationships that the neural architecture effectively leverages. Conversely, some models highlight unique features not prominently featured in others. For instance, the Neural Network’s *ECIndex* does not appear in other models’ top ten, suggesting its specific relevance within the neural network’s processing framework. Similarly, SVM uniquely identifies *VSA_EState4* and *nBase* as important features, potentially indicating their specialized role in the high-dimensional feature space managed by the support vector algorithm.

From the biological context, the linear models (SVM and logistic regression) revealed distinct feature contributions to classification outcomes, with bidirectional coefficients highlighting structure-activity relationships (Figure 2). In the SVM model, *EState_VSA9* (0.00915) and *BIC4* (0.00866) enhanced the positive class outcome, suggesting hydrogen-bonded surface regions and branched molecular connectivity improving target engagement through hydrophilic interactions or multi-site binding. Conversely, *VSA_EState4* (-0.00963) and *nBase* (-0.00938) exhibited negative correlations, implying excessive hydrophobicity or basic group density may hinder solubility or membrane permeability. Similarly, in logistic regression, *MATS1c* (0.02438) and *AATS2v* (0.02409) positively influenced predictions, reflecting the importance of charge distribution patterns and steric volume alignment for electrostatic and shape complementarity. In contrast, *GATS3dv* (-0.02495) and *GATS3pe* (- 0.02466) reduced activity, likely due to uniform electron distribution or excessive polarizability destabilizing ligand-receptor complexes. These descriptors collectively encode electronic, steric, and topological properties critical to drug-receptor binding and pharmacokinetics, aligning with established principles of molecular recognition and bioavailability[57–59].

Furthermore, excipients such as glycine or dextrose, often considered inert, have been shown to influence the stability and promote growth of the probiotic bacteria in the study. For instance, descriptors like *AATSC1se* and *AATSC1pe*, which are prominent across multiple models such as Random Forest and Gradient Boosting, respectively are likely to capture molecular attributes related to electronic or steric effects that directly impact excipient functionality. These features may correlate with critical roles such as stabilizing active pharmaceutical ingredients (APIs), maintaining tonicity, or preventing degradation during storage and administration. Additionally, descriptors like *ETA_dPsi_A* and *NsCl* emerge as significant contributors to the predictive models (Random Forest and Gradient Boosting, respectively), offering deeper insights into excipient behaviour. *ETA_dPsi_A* reflects hydrogen bonding propensity, which is crucial for interactions between excipients and APIs, influencing drug stability, solubility, and bioavailability. *NsCl* represents the count of chlorine atoms in a molecule and highlights the importance of chemical reactivity and potential toxicity considerations in excipient selection.

However, the emphasis of the neural network model on ATS descriptors (e.g., *ATS8Z*, *ATS2m*, *ATS0v*) and *ECIndex* highlights the distribution of specific molecular properties across a molecule’s topology. These descriptors are calculated by correlating atomic properties, such as polarizability, mass, or van der Waals volume, over defined topological distances within the molecule. The biological significance of these descriptors lies in their ability to quantify structural and physicochemical attributes that influence molecular interactions with biological systems. For instance, polarizability (captured by *ATS8Z*) is crucial for understanding intermolecular forces such as hydrogen bonding and van der Waals interactions, which play a key role in drug-receptor binding or excipient-API stabilization. Atomic mass distribution (*ATS2m*) can correlate with molecular stability and transport properties, while van der Waals volume (*ATS0v*) provides insights into steric hindrance and molecular fit within biological targets. Together, these molecular descriptors provide a comprehensive understanding of the structural and physicochemical attributes that govern excipient performance in stabilizing the probiotics being studied. The aforementioned strategy underscores the nuanced balance required in molecular design to optimize both biological activity and pharmacologically desirable properties.

This approach underscores the relevance of leveraging multiple models to capture different aspects of feature importance. Compiling insights from diverse models allow for a more holistic understanding of the dataset, enhancing the robustness of feature selection and its interpretation.

#### 3.5. Active Machine Learning Results

To systematically evaluate the effects of excipients on *Lactobacillus plantarum* growth, five distinct machine learning models were applied: Neural Networks, Gradient Boosting, Logistic Regression, Support Vector Machines (SVM), and Random Forest. Each model was trained and refined over three rounds of active learning, with certainty and error metrics recorded at each stage to assess predictive stability and accuracy.

The performance of these models varied significantly, with the Neural Network demonstrating the highest certainty, reaching 88.84% in the final round. However, it is noteworthy that the model also exhibited a relatively high error rate of 16.77%. This suggests that while Neural Networks excel in identifying potential patterns, they may benefit from further refinement to reduce error. Gradient Boosting followed closely, achieving a final certainty of 87.57% and the lowest error rate among all models at 14.57%, indicating strong stability and a favourable balance between certainty and error. Logistic Regression also showed robust performance, attaining 85.33% certainty with a 15.93% error rate, making it a reliable alternative for capturing excipient-probiotic interactions.

By contrast, Random Forest and SVM demonstrated lower final certainty levels, with Random Forest reaching 51.54% certainty and a 4.92% error rate, and SVM attaining 51.88% certainty with a 6.45% error rate. The comparatively low certainty in these models indicates the limitation simpler algorithms come across when handling complex, non-linear relationships within the excipient-probiotic dataset, highlighting the need for more advanced or ensemble approaches.

The increase in model certainty over successive active learning rounds revealed unique improvement patterns for each approach. Neural Networks showed a steady increase, with certainty values rising from 86.55% in the initial round to 88.84% in the final round. Gradient Boosting followed a similar trajectory, progressing consistently from an initial certainty of 84.81% to 87.57% by the last round. Logistic Regression, while starting at a lower initial certainty (74.85%), showed increase in certainty levels at 85.26% by the final iteration. In contrast, SVM showed fluctuations in model certainty, dropping from 53.44% in the first round to 41.09% in the second before increasing to 51.88% in the final round. Random Forest also showed similar patterns. These trends highlight the ability of more complex models, particularly Neural Networks and Gradient Boosting, to sustain and build upon predictive certainty through iterative learning, while simpler models such as SVM and Random Forest displayed less stability in their certainty improvements.

These results illustrate that more sophisticated models, particularly Neural Networks and Gradient Boosting, showed consistent improvement in certainty across rounds, underscoring their capacity to capture intricate patterns in excipient-bacterial interactions. Logistic Regression, while simpler, also demonstrated strong progressive gains, indicating its adaptability to structured active learning. The Random Forest and SVM models showed fluctuating performance, which may reflect limitations in their ability to generalize from small datasets with complex interactions.

Overall, the active learning approach enabled targeted refinement of model predictions, particularly benefiting the more complex models. These findings suggest that incorporating advanced models, such as Neural Networks and Gradient Boosting, in an active learning framework can significantly enhance predictive reliability and accuracy, facilitating improved excipient selection for precision probiotic applications.

#### 3.6. Model Validation

To assess the predictive performance and reliability of the machine learning models, a model validation step was conducted using two specific excipients: Sodium Chloride and Potassium Bicarbonate. Both excipients were categorized as having a neutral effect on the growth of *Lactobacillus plantarum*, serving as critical benchmarks to evaluate the models’ accuracy and confidence in their predictions.

Random Forest predicted both Sodium Chloride and Potassium Bicarbonate as neutral with certainty levels of 46.0% and 52.0%, respectively. These relatively modest confidence scores suggest that while the Random Forest model correctly identified the neutral classification, there is room for improvement in terms of model confidence.

Support Vector Machine (SVM) model demonstrated slightly higher certainty levels, predicting both excipients as neutral with certainties of 54.65% for Sodium chloride and 56.91% for Potassium bicarbonate. Although these certainty values are higher than those of Random Forest, they still reflect a moderate level of confidence, highlighting potential limitations in the SVM’s ability to decisively classify these samples.

Logistic Regression exhibited markedly high certainty in its predictions, assigning a 99.99% certainty to Sodium chloride and a 99.96% certainty to Potassium bicarbonate, correctly classifying as neutral on both occasions. These exceptionally high confidence scores indicate that the Logistic Regression model is extremely confident about its neutral predictions, suggesting strong model reliability for these specific cases, and possibly for excipients with similar molecular descriptors.

The Gradient Boosting model also showed robust performance, with a certainty of 89.27% for Sodium chloride and an impressive 99.88% for Potassium bicarbonate, both correctly identified as neutral. The high certainty associated with Potassium bicarbonate aligns closely with the Logistic Regression model, underscoring Gradient Boosting’s effectiveness in confidently classifying neutral excipients.

Lastly, the Neural Network model achieved high certainty levels, predicting both excipients as neutral with 99.14% certainty for Sodium chloride and 99.54% certainty for Potassium bicarbonate. These results are comparable to those of Logistic Regression and Gradient Boosting, reflecting the Neural Network’s strong confidence and reliability in its predictions.

Overall, the validation results indicate that while all models successfully classified the two excipients as neutral, there is significant variability in their confidence levels. Logistic Regression, Gradient Boosting, and Neural Networks consistently demonstrated high certainty in their predictions, suggesting superior reliability and robustness in handling these specific cases. In contrast, Random Forest and SVM exhibited lower certainty levels, highlighting potential areas for model refinement to enhance predictive confidence. These findings further underline the significance of selecting appropriate models based on their confidence and reliability, especially when making critical predictions in active learning frameworks.

## 4. Conclusions

This study demonstrates the capability of active machine learning to accurately predict the effects of pharmaceutical excipients on the growth of the probiotic *Lactobacillus plantarum*. Initiating the process with a dataset of just nine excipient-bacteria interactions, the active learning framework was employed to extend predictions to an additional 116 untested excipients. Five distinct machine learning models—Neural Networks, Gradient Boosting, Logistic Regression, Support Vector Machines (SVM), and Random Forest were systematically trained and refined over three rounds of active learning, each iteration enhancing the models’ predictive capabilities.

The active learning approach produced varied outcomes across the models. Neural Networks demonstrated strong proficiency in identifying complex patterns, emerging as the most effective model in this study. Gradient Boosting followed closely, showcasing stability and balanced predictive performance. Logistic Regression also proved to be a reliable option, effectively capturing excipient-probiotic interactions. These results underscore the ability of different algorithms to address the challenges of modeling such interactions, while highlighting the advantages of more advanced techniques in achieving consistent performance.

Conversely, Random Forest and SVM models exhibited lower final certainty levels, with Random Forest achieving 51.54% certainty and an error rate of 4.92%, and SVM attaining 51.88% certainty with a 6.45% error rate. These results suggest that while simpler algorithms like Random Forest and SVM are capable of correctly classifying neutral excipients, their moderate certainty levels highlight potential limitations in capturing the intricate patterns and dependencies within the dataset. This defines the necessity for advanced or ensemble-based approaches to improve predictive confidence and reliability.

Feature importance analysis identified critical molecular descriptors, including the eccentric connectivity index (ECIndex) and various atomic triplet descriptors (e.g., GATS3dv, AATS1se), as pivotal in determining excipient effects. These descriptors align with existing literature, emphasizing the significance of detailed molecular features in enhancing machine learning model performance for biological activity prediction. Additionally, Principal Component Analysis (PCA) provided valuable insights into the data distribution’s evolution throughout the active learning process, illustrating the algorithm’s effectiveness in capturing a diverse and informative sample space essential for model generalization.

The findings of this research highlight active machine learning’s efficacy in optimizing predictive models for pharmaceutical sciences, particularly when working with limited datasets. By iteratively refining model predictions and identifying key molecular features, active ML facilitates informed excipient selection, thereby advancing the development of precision probiotics tailored for optimal therapeutic outcomes. Our work could motivate expanding the dataset to encompass a broader array of excipients and exploring advanced feature representation techniques, especially for polymeric excipients whose properties significantly influence interactions with probiotics.

## Supporting information

Train_Initial

Train_Final

Test_Initial

## Author Contributions

**A.P.**, **M.A.** and **S.S.N.** are equally contributing authors. **A.P.**, Data curation; Formal analysis; Investigation; Methodology; Resources; Software; Validation; Visualization; Writing – original draft; Writing – review & editing. **M.A.**, Data curation; Formal analysis; Funding acquisition; Investigation; Methodology; Resources; Validation; Visualization; Writing – original draft; Writing – review & editing. **S.S.N.**, Data curation; Formal analysis; Investigation; Methodology; Resources; Software; Validation; Visualization; Writing – original draft; Writing – review & editing. **A.K.N.**, Data curation; Methodology; Investigation; Software; Validation; Visualization; Writing – original draft; Writing – review & editing. **S.K.N.**, Data curation; Formal analysis; Methodology; Resources; Supervision; Validation; Writing – review & editing. **S.K.D.**, Data curation; Formal analysis; Methodology; Resources; Writing – review & editing. **S.N.**, Conceptualization; Formal analysis; Funding acquisition; Investigation; Methodology; Project administration; Resources; Supervision; Validation; Writing – original draft; Writing – review & editing.

## Acknowledgement

This work and S.S.N were supported by DD Innovations Incorporated, U.S.A. (Ref No. DD/KIIT/001/2023). M.A. was supported by CSIR University Grants Commission (UGC) Junior Research Fellowship Program [Ref No. 221610080088/(CSIR- UGC NET OCT. 2022)].

## Data availability statement

All data that support the findings of this study are included within the article and supplementary materials.

