## Supplementary Materials for "Enhanced Formulation of Precision Probiotics through Active Machine Learning"

### *Table of Contents*

*Growth Curves of Lactobacillus plantarum in presence and absence of different pharmaceutical grade excipients.* .......... *S3*

*Support Vector Machines (SVM)* .......... *S8*

*Detailed Explanation of Key Concepts* .......... *S8*

*Hyperparameter Tuning* .......... *S9*

*Model Performance Metrics* .......... *S9*

*Logistic Regression* .......... *S10*

*Detailed Explanation of Key Concepts* .......... *S10*

*Hyperparameter Tuning* .......... *S11*

*Model Performance Metrics* .......... *S11*

*Gradient Boosting* .......... *S12*

*Detailed Explanation of Key Concepts* .......... *S12*

*Hyperparameter Tuning* .......... *S13*

*Model Performance Metrics* .......... *S13*

*Neural Network Architecture* .......... *S14*

*Key Concepts in Neural Networks* .......... *S15*

*Custom KerasWrapper Class and Integration with Active Learning* .......... *S15*

**Growth Curves of Lactobacillus plantarum in presence and absence of different pharmaceutical grade excipients.**

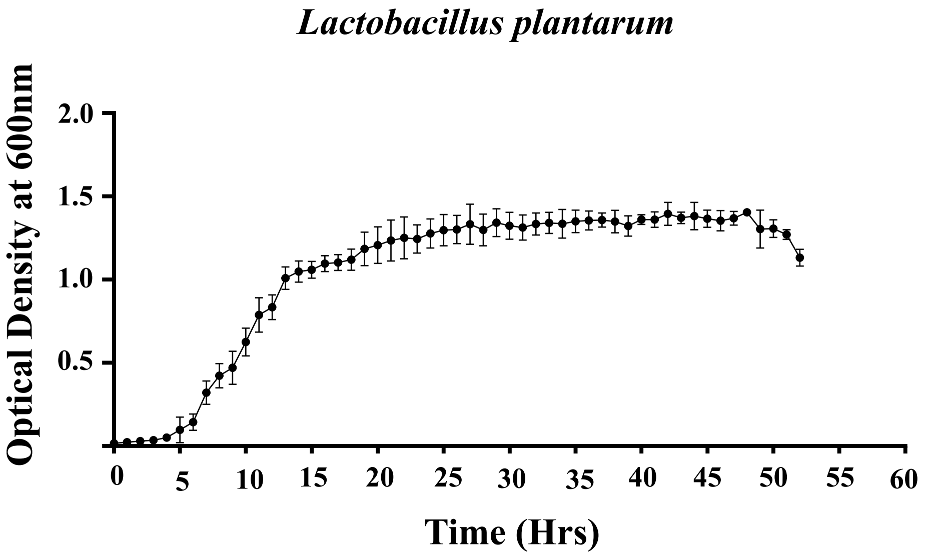


**Figure S1.** Growth curve pattern of *Lactobacillus plantarum* over a period of 48 hours in MRS broth. Error bars represent ±1 standard error of mean (SEM) from mean for three (n=3) independent sets of experiments.

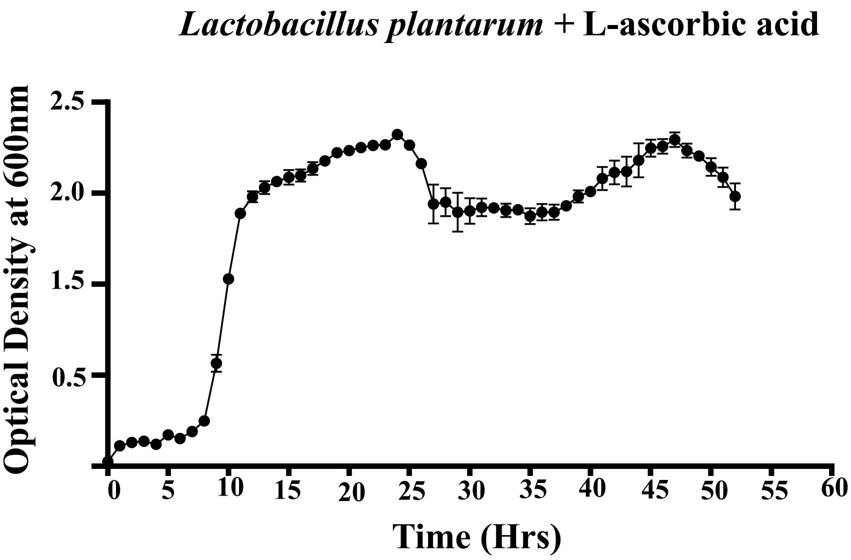


**Figure S2.** Growth curve pattern of *Lactobacillus plantarum* supplemented with L-ascorbic acid at a concentration of 70 μM over a period of 48 hours in MRS broth. Error bars represent ±1 standard error of mean (SEM) from mean for three (n=3) independent sets of experiments.

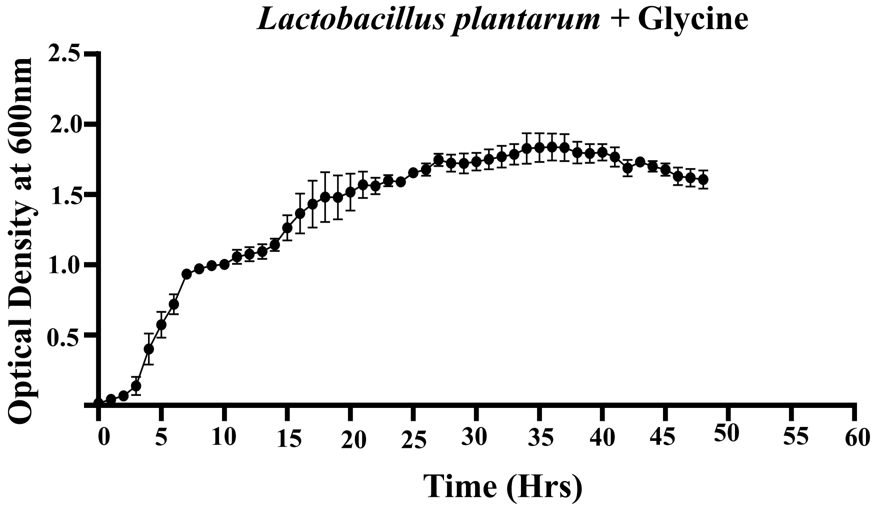


**Figure S3.** Growth curve pattern of *Lactobacillus plantarum* supplemented with Glycine at a concentration of 70 µM over a period of 48 hours in MRS broth. Error bars represent ±1 standard error of mean (SEM) from mean for three (n=3) independent sets of experiments.

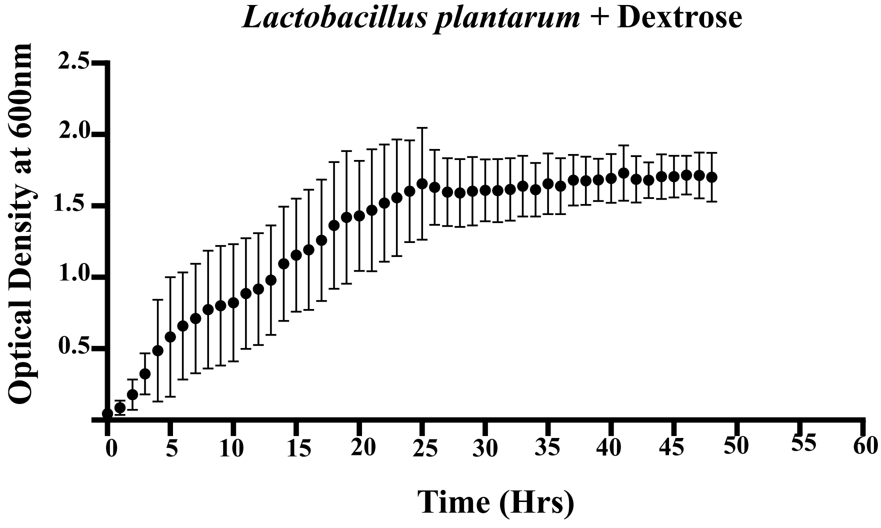


**Figure S4.** Growth curve pattern of *Lactobacillus plantarum* supplemented with Dextrose at a concentration of 70 µM over a period of 48 hours in MRS broth. Error bars represent ±1 standard error of mean (SEM) from mean for three (n=3) independent sets of experiments.

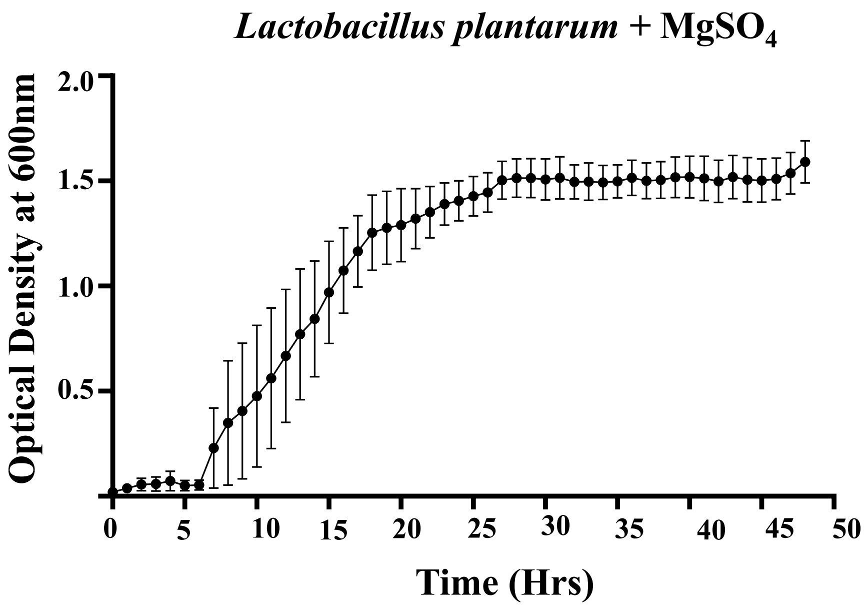


**Figure S5.** Growth curve pattern of *Lactobacillus plantarum* supplemented with Magnesium Sulphate ($MgSO_4$) at a concentration of 70 μM over a period of 48 hours in MRS broth. Error bars represent ±1 standard error of mean (SEM) from mean for three (n=3) independent sets of experiments.

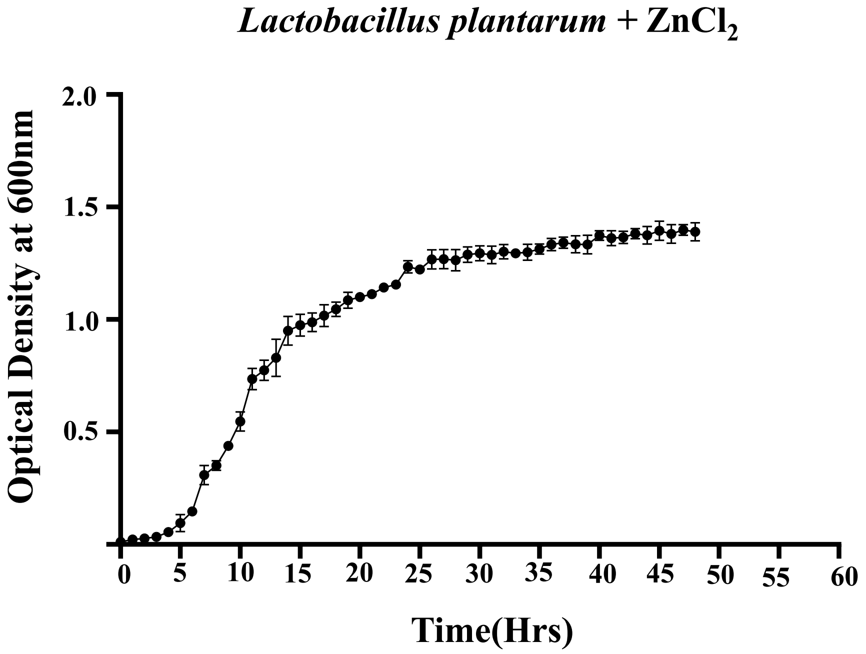


**Figure S6.** Growth curve pattern of *Lactobacillus plantarum* supplemented with Zinc Chloride ($ZnCl_2$) at a concentration of 70 μM over a period of 48 hours in MRS broth. Error bars represent ±1 standard error of mean (SEM) from mean for three (n=3) independent sets of experiments.

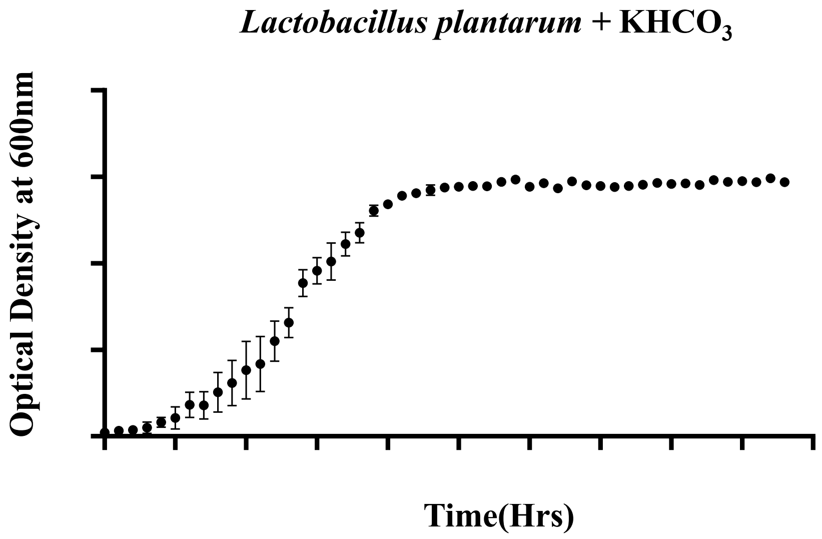


**Figure S7.** Growth curve pattern of *Lactobacillus plantarum* supplemented with Potassium Bicarbonate ($KHCO_3$) at a concentration of 70 μM over a period of 48 hours in MRS broth. Error bars represent ±1 standard error of mean (SEM) from mean for three (n=3) independent sets of experiments.

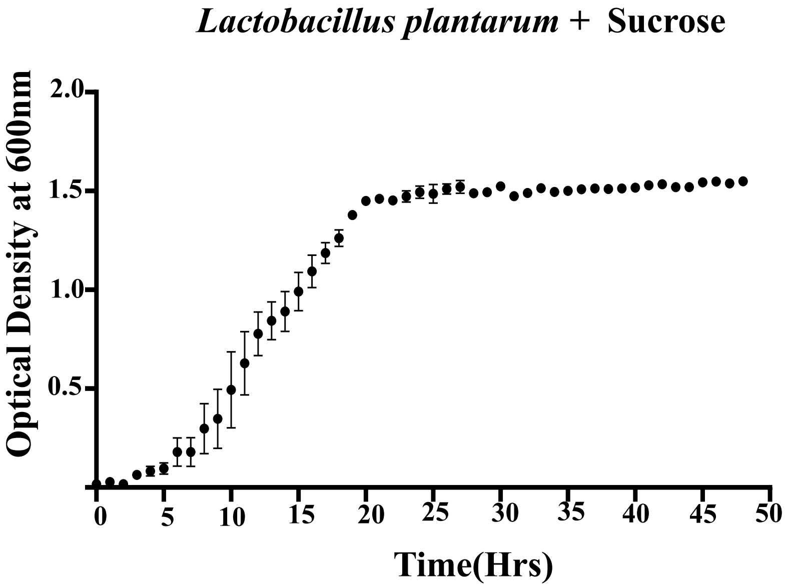


**Figure S8.** Growth curve pattern of *Lactobacillus plantarum* supplemented with Sucrose at a concentration of 70 μM over a period of 48 hours in MRS broth. Error bars represent ±1 standard error of mean (SEM) from mean for three (n=3) independent sets of experiments.

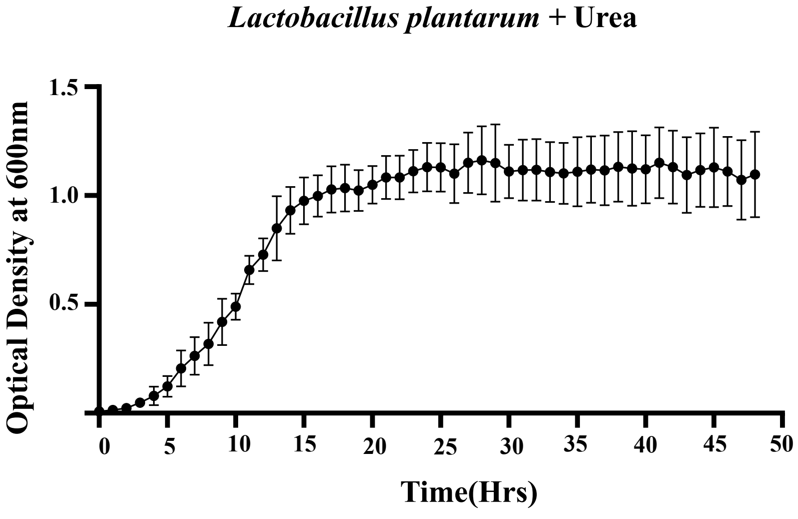


**Figure S9.** Growth curve pattern of *Lactobacillus plantarum* supplemented with Urea at a concentration of 70 µM over a period of 48 hours in MRS broth. Error bars represent ±1 standard error of mean (SEM) from mean for three (n=3) independent sets of experiments.

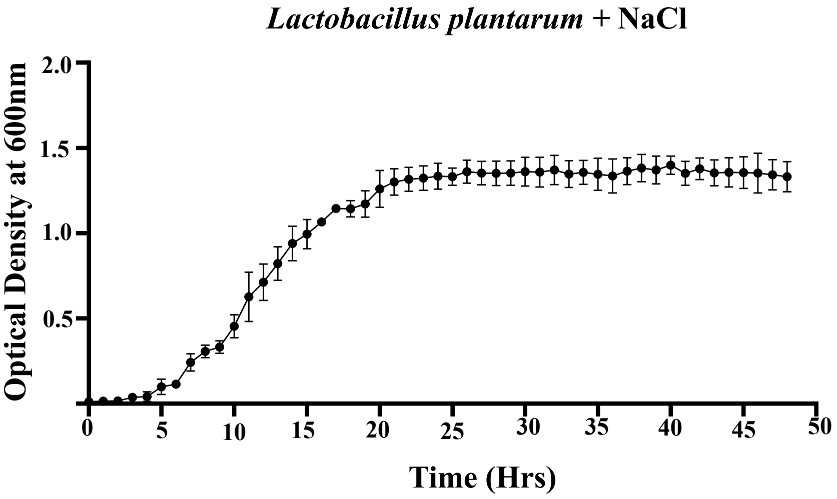


**Figure S10.** Growth curve pattern of *Lactobacillus plantarum* supplemented with Sodium Chloride (NaCl) at a concentration of 70 µM over a period of 48 hours in MRS broth. Error bars represent ±1 standard error of mean (SEM) from mean for three (n=3) independent sets of experiments.

**Support Vector Machines (SVM)**

Support Vector Machines are a class of supervised learning algorithms that aim to find the optimal hyperplane that separates data points into distinct classes. The hyperplane is a decision boundary defined in the feature space, which may be high-dimensional or even infinite-dimensional depending on the choice of kernel. The margin of the hyperplane is defined as the distance to the closest data points, known as support vectors. A larger margin reduces overfitting and improves generalization. In this study, a linear kernel was selected to address the linear separability of the dataset.

Detailed Explanation of Key Concepts

1. **Hyperplane:**

   A hyperplane is a geometric concept representing a decision boundary that divides the feature space into distinct regions, each corresponding to a class label. For instance, in a two-dimensional feature space, the hyperplane is a line that separates two classes. In higher-dimensional spaces, it becomes a plane or hyperplane.

2. **Margin and Support Vectors:**

   The margin refers to the distance between the hyperplane and the nearest data points from either class, known as support vectors. The SVM algorithm seeks to maximize this margin to create a decision boundary that generalizes well to unseen data. A larger margin typically reduces the risk of overfitting, improving the model's ability to classify new data points accurately.

3. **Linear Kernel:**

   A linear kernel is a mathematical function used in SVMs to compute the similarity between data points. It assumes that the classes are linearly separable, meaning that a straight line (or flat hyperplane) can divide them in the feature space. This kernel is computationally efficient and effective when the dataset does not require complex, non-linear boundaries for separation.

4. **Linear Separability:**

   Linear separability means that the classes in a dataset can be separated by a straight line (in 2D) or a flat hyperplane (in higher dimensions). When the dataset exhibits this property, the SVM with a linear kernel can effectively classify the data with a simple decision boundary.

5. **Probability Estimates for Uncertainty Sampling:**
   Enabling probability estimates in the SVM model allows it to compute probabilities for each class prediction. These probabilities provide a measure of confidence in the prediction. In active learning, uncertainty sampling leverages these probabilities to identify the data points for which the model has the least confidence, ensuring that the most ambiguous and informative samples are prioritized for labelling.
6. **Decision Boundary:**
   The decision boundary is the hyperplane that separates different classes in the feature space. It defines the point at which the model transitions from predicting one class to another. The location and orientation of this boundary are determined by the model parameters and the training data.

Hyperparameter Tuning

No hyperparameter tuning was conducted for the SVM model in this study. Instead, the default parameters of the scikit-learn SVC class were used. The decision to use default parameters was made because the model's performance within the active learning framework was sufficient to meet the study's objectives.

Model Performance Metrics

The performance of the SVM was evaluated using model certainty and error bars. These metrics provided insights into the model's robustness and ability to generalize, especially when integrated into the active learning framework.

1. Model Certainty: This metric reflects the model's confidence in its predictions across the dataset.
2. Error Bars: These illustrate variability in predictions, highlighting areas of potential improvement or uncertainty.

Unlike traditional metrics such as accuracy, precision, recall, and F1 score, which offer static evaluations, uncertainty-based metrics provide a dynamic and iterative perspective, better suited to active learning methodologies.

**Logistic Regression**

Logistic Regression is a linear model used for binary classification tasks, where the probabilities of class membership are modelled using a logistic function. The model outputs a probability score between 0 and 1, which is thresholded to classify data points into one of two classes. Logistic Regression is a straightforward yet effective algorithm, particularly suited for datasets where the relationship between features and the target variable is approximately linear.

Detailed Explanation of Key Concepts

1. **Logistic Function:**

   The logistic function, also known as the sigmoid function, is defined as:

   $$f(x) = \frac{1}{1 + e^{-x}}$$

   This function maps any real-valued number into the range [0, 1], making it ideal for modelling probabilities. In Logistic Regression, the function is applied to the weighted sum of input features to compute the probability of class membership.

2. **Decision Boundary:**

   Logistic Regression assumes a linear relationship between the features and the log-odds of the target variable. The decision boundary is a hyperplane in the feature space that separates the two classes, defined by the equation:

   $$w_1x_1 + w_2x_2 + \cdots + w_nx_n + b = 0$$

   Here, $w$ represents the weights, $x$ the features, and $b$ the bias term. Data points on one side of the boundary are classified as one class, while points on the other side belong to the opposite class.

3. **Probabilistic Predictions**:

   Logistic Regression outputs probabilistic predictions, where the binary classification decision is made on the basis of a prediction probability threshold (typically chosen as 0.5) These probabilities also provide a measure of confidence in the model's predictions, which is critical for uncertainty sampling in active learning.

4. **Uncertainty Sampling:**

   In the active learning framework, the Logistic Regression model employed uncertainty sampling to prioritize data points for labelling. The model focused on instances where the predicted probability was closest to 0.5, indicating high uncertainty. By iteratively labelling and training on these ambiguous samples, the model progressively improved its decision boundary and classification accuracy.

##### Hyperparameter Tuning

No hyperparameter tuning was conducted for Logistic Regression in this study. The default parameters of the LogisticRegression class from Python's scikit-learn library were used, as they provided sufficient performance within the active learning framework. Regularization strength ($C$) and solver were left at their default values, and further optimization was deemed unnecessary given the model's straightforward nature and satisfactory results.

##### Model Performance Metrics

The performance of the Logistic Regression model was evaluated using uncertainty-based metrics rather than traditional measures such as accuracy, precision, recall, or F1 score.

1. Model Certainty: This metric quantified the confidence of the model's predictions over the entire dataset, providing insights into how certain the model was about its classifications.
2. Error Bars: These metrics were used to visualize the variability in the model's predictions across successive active learning cycles, highlighting areas where performance could be improved.

The emphasis on uncertainty-based metrics aligned with the iterative and dynamic nature of active learning, where the goal is to improve model performance through informed sampling of uncertain data points.

**Gradient Boosting**

Gradient Boosting is an ensemble method that combines the predictions of weak learners (typically shallow decision trees) to form a stronger predictive model. The key features of Gradient Boosting are its stage-wise construction and the use of a loss function to iteratively minimize prediction errors. At each stage, the algorithm fits a new decision tree to the negative gradient of the loss function, effectively correcting errors made by the previous ensemble.

Detailed Explanation of Key Concepts

1. **Decision Tree:**
   A decision tree is a machine learning model that represents decisions and their possible consequences as a tree-like structure. It splits the dataset into subsets based on feature values, with each split representing a decision that leads to one of several outcomes. In a classification task, the leaves of the tree correspond to class labels. Decision trees are easy to interpret and computationally efficient, making them well-suited as weak learners in Gradient Boosting.
2. **Loss Function:**
   The loss function measures the discrepancy between predicted and actual values, serving as a guide for model optimization. In classification tasks, commonly used loss functions include log-loss (cross-entropy), which penalizes incorrect classifications based on their predicted probabilities. Gradient Boosting relies on the differentiable nature of the loss function to compute gradients, enabling precise updates to the model at each stage.
3. **Differentiable Loss Function:**
   A differentiable loss function is one that can be mathematically differentiated, meaning that its gradient can be calculated. This property is crucial for Gradient Boosting, as the algorithm uses the gradient of the loss function to iteratively adjust the predictions of its weak learners, reducing errors in a systematic manner.
4. **Weak Learners (Decision Trees):**
   Gradient Boosting typically employs shallow decision trees as weak learners. These trees are computationally efficient and capture local patterns in the data, making them ideal for ensemble methods.
5. **Uncertainty Sampling in Active Learning:**
   The active learning framework integrated Gradient Boosting with uncertainty sampling to prioritize samples for labeling. The model focused on predictions with the highest

uncertainty, ensuring that labeling efforts were directed at data points most likely to improve the ensemble's decision boundaries.

Hyperparameter Tuning

No hyperparameter tuning was conducted for Gradient Boosting in this study. The default settings of the GradientBoostingClassifier class from Python’s scikit-learn library were used. Parameters such as the number of estimators, learning rate, and maximum depth of individual trees were left unchanged, as the default configuration provided adequate performance for the task.

Model Performance Metrics

The performance of the Gradient Boosting model was evaluated using uncertainty-based metrics tailored to the iterative nature of active learning:

1. Model Certainty: Quantified the confidence of the model's predictions across the dataset, providing insights into its reliability.
2. Error Bars: Visualized the variability in predictions, highlighting areas where the model's performance could be further refined.

These metrics offered a dynamic evaluation framework, emphasizing the progressive improvement of the model during active learning cycles, rather than static assessments like accuracy, precision, or F1 score.

**Neural Network Architecture**

1. Layers in a Neural Network:

   A neural network is composed of layers, each consisting of artificial neurons that process and pass data to the next layer.

   - Dense Layer (Fully Connected): Each neuron in the Dense layer is connected to every neuron in the previous layer, enabling the network to learn complex patterns.
   - Dropout Layer: Dropout is a regularization technique that randomly disables a fraction of neurons during training to reduce overfitting and improve generalization.

2. Output Layer with Softmax Activation:

   The output layer contains three neurons, corresponding to the three target classes in the dataset. The softmax activation function converts the raw outputs into probabilities, ensuring that the sum of probabilities across all classes equals 1. This makes it suitable for multi-class classification tasks.

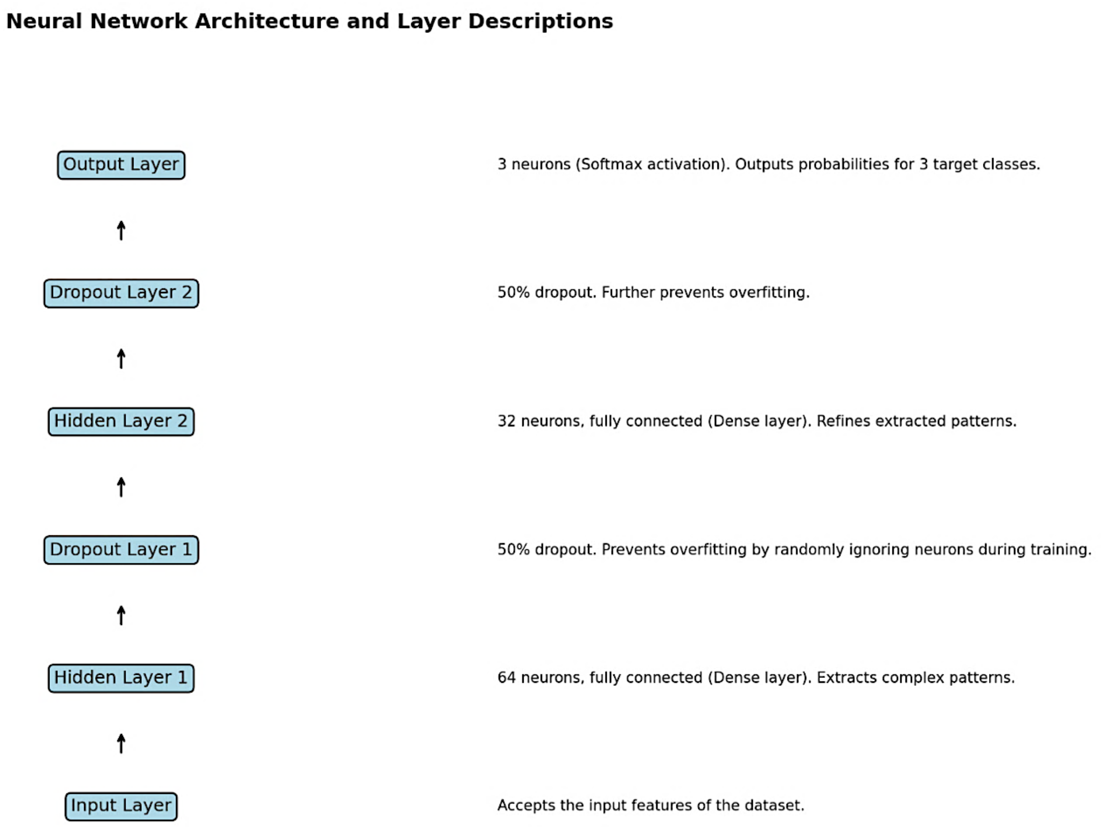

Key Concepts in Neural Networks

1. Activation Functions:
   Activation functions introduce non-linearity to the model, enabling it to learn complex relationships. The softmax activation function in the output layer ensures that the model predicts probabilities for each class.
2. Optimizer (Adam):
   The optimizer adjusts the weights of the neurons to minimize the loss function. Adam is an adaptive optimizer that combines the benefits of AdaGrad and RMSProp, making it efficient for deep learning tasks.
3. Loss Function (Categorical Cross-Entropy):
   The loss function measures the error between the predicted probabilities and the actual class labels. Categorical cross-entropy is commonly used for multi-class classification tasks and encourages the model to output high probabilities for the correct class.
4. Epochs and Batch Size:
   - Epochs: Represent the number of complete passes through the training dataset. In this study, 50 epochs allowed the model to learn effectively.
   - Batch Size: Refers to the number of samples processed before updating the model's weights. A batch size of 32 balanced computational efficiency and learning precision.

**Custom KerasWrapper Class and Integration with Active Learning**

A custom KerasWrapper class was implemented to bridge the neural network with scikit-learn's ActiveLearner. This wrapper provided methods for:

- Model Training: Fitting the model to the labelled dataset.
- Prediction: Generating class probabilities for unlabelled samples.
- Uncertainty Sampling: Identifying data points with the highest uncertainty for querying and labeling.

This integration enabled the iterative improvement of the neural network through the active learning pipeline.
